# Olfactory Sensitivity Aligns with Caste Revealing Multipotential Minors and Specialized Soldier Majors in *Camponotus floridanus* Ants

**DOI:** 10.1101/2022.04.01.486728

**Authors:** S.T. Ferguson, I. Bakis, N.D. Edwards, L.J. Zwiebel

## Abstract

*Camponotus floridanus* ant colonies are comprised of a single reproductive queen and thousands of sterile female offspring that consist of two morphologically distinct castes: smaller minors and larger majors. Minors perform most of the tasks within the colony, including brood care and food collection, whereas majors have fewer clear roles and have been hypothesized to act as a specialized solider caste associated with colony defense. The allocation of workers to these different tasks depends on the detection and processing of local information including pheromones and other chemical blends such as cuticular hydrocarbons. We examined the electrophysiological responses to general odorants, cuticular extracts, and a trail pheromone in adult minor and major *C. floridanus* workers, revealing that the repertoire of social behaviors is positively correlated with olfactory sensitivity. Minors in particular display primarily excitatory responses to olfactory stimuli, whereas major workers respond primarily with inhibitory signals. The notable exception to this paradigm is that both minors and majors display robust, dose-dependent excitatory responses to conspecific, non-nestmate cuticular extracts. Moreover, while both minors and majors actively aggress non-nestmate foes, majors display significantly enhanced capabilities to rapidly subdue and kill opponents. Overall, our data suggest that *C. floridanus* majors do indeed represent a physiologically and behaviorally specialized soldier caste and support a model in which caste-specific olfactory sensitivity plays an important role in task allocation and the regulation of social behavior in ant colonies.

**Significance Statement:** The detection and odor coding of chemical cues are essential components of the collective behavior observed in eusocial ants. To better understand the interdependent relationship between olfactory sensitivity and the allocation of worker castes to the various tasks critical for the success of the colony, a series of behavioral assessments and an electrophysiological survey of the antennae comprising general odorants, cuticular extracts, and a trail pheromone were undertaken. These studies reveal the behavioral repertoire of minors and majors aligned with profound shifts in olfactory sensitivity and odor coding. Our data support the hypothesis that minors are multipotential workers with broad excitatory sensitivity, and majors are dedicated soldiers with a highly specialized olfactory system for the detection of non-nestmate foes.

## Introduction

In ants colony performance and survival depend on a dynamic and decentralized process of distributing work across all members of the colony (Gordon 1996). Collective behaviors such as nursing offspring, foraging for food, and nest defense emerge as groups of ants detect and respond to a broad range of local information including, most notably, pheromones and other odors such as cuticular hydrocarbons (CHCs) (Hölldobler and Wilson 1990). In addition, the colonies of certain ant genera contain morphologically distinct worker castes that may perform specialized roles within the colony. The behavioral repertoire of army ants (genus *Eciton*), for example, is associated with worker size and shape (Powell and Franks 2006; Franks, Sendova-Franks, and Anderson 2001). Here, the largest workers form a soldier caste that performs a restricted subset of behaviors primarily specialized for nest defense. In addition to their distinctive sharply pointed and sickle-shaped mandibles, the brain volume of these soldiers, including the antennal lobe (AL) and mushroom bodies (MB), are significantly reduced compared with other workers within the colony, suggesting that adaptive changes in these olfactory processing centers of the brain may contribute to or otherwise reflect differences in worker behavior (O’Donnell et al. 2018). Similarly, in carpenter ants (genus *Camponotus*), smaller minor workers carry out the majority of tasks within the colony and correspondingly have larger AL volumes and more AL glomeruli than their larger major worker sisters (Zube and Rössler 2008; Zube et al. 2008). Transcriptome profiling in *C. floridanus* studies reveals that mRNA transcripts associated with muscle development are enriched in majors, whereas transcripts associated with synaptic transmission, cell-cell signaling, neurological system processes, and behavior are upregulated in the smaller but more behaviorally diverse minors (Simola et al. 2013). Taken together, these anatomical and molecular changes have given rise to colloquial characterizations of “brainy” minors and “brawny” majors in these species and and suggest that differences in olfactory processes may contribute to the specialized behavioral repertoire of subsets of workers.

Numerous behavioral and chemical ecological studies as well as recent advances in targeted molecular approaches and gene editing techniques highlight the importance of the chemical senses in driving social behaviors in ants and other Hymenoptera (reviewed in (Ferguson, Anand, and Zwiebel 2020) and (Ferguson, Bakis, and Zwiebel 2021)). In ants and other insects, the peripheral detection of pheromones and other chemical signals occurs principally in olfactory sensory neurons (OSNs) in the antennae and other sensory appendages. OSNs express chemoreceptor gene families that include odorant receptors (ORs), ionotropic receptors (IRs) and gustatory receptors (GRs) (Ferguson, Bakis, and Zwiebel 2021). In ants, genomic studies revealed that the OR gene family has rapidly evolved through a birth-and-death process that resulted in a large expansion of these gene such that ants have the largest number of ORs among any insect species. (Zhou et al. 2012; Zhou et al. 2015). In alignment with the hypothesis that the large expansion of chemoreceptor gene families facilitated the evolution of eusocial insect colonies, it has been demonstrated that ORs are act in the detection of general odorants and critical social cues, including CHCs (Slone et al. 2017; Pask et al. 2017). Gene editing in two primitively eusocial ants to generate null mutations of the obligate OR co-receptor (Orco) gene, which is required for the formation and functionality of odor-gated OR ion channels, resulted in significant deficits in odor sensitivity as well as reduced colony cohesion and severe alterations in the AL development (Yan et al. 2017; Trible et al. 2017). Furthermore, targeted pharmacological modulation of Orco function in wildtype *C. floridanus* workers with unaffected AL and central brains significantly impaired aggression-mediated non-nestmate recognition (Ferguson et al. 2020). Taken together, these studies highlight the critical role that olfaction plays in the detection of pheromones and other behaviorally salient odorants that ultimately act to mediate social behaviors in ants. However, considerably less is known about modulation of peripheral olfactory sensitivity to pheromones and other general odorants between morphological castes.

An appreciation as to how variation in olfactory signaling contributes to or, alternatively, reflects the unique roles performed by different worker castes within the colony will better inform our understanding of the emergent properties of coordinated social behavior in ants and other eusocial animals. To begin to address that question, we have undertaken a behavioral assessment as well as a broad electrophysiological survey of the antennae across more than 400 general odorants, non-nestmate cuticular extracts, and a known trail pheromone in both unitary and blended stimulation paradigms across the two morphological *C. floridanus* worker castes. These studies examine the hypothesis that alterations in peripheral olfactory sensitivity correlate with the distinctive social behaviors of these morphological castes. Our data confirm that the majority of tasks in *C. floridanus* colonies are indeed carried out by minor workers and that majors have a more specialized defensive role within the colony. In keeping with their more diverse task engagement, peripheral responses to general odor blends and trail pheromone were significantly higher in minors than majors. Moreover, minor workers primarily displayed a broad range of excitatory responses while majors manifested primarily inhibitory responses suggesting a fundamental shift in odor coding between these castes. In contrast, CHCs and other hydrophobic compounds associated with the *C. floridanus* cuticle elicited robust excitatory responses from both majors and minors. Behaviorally, majors were significantly more adept at rapidly subduing opponent non-nestmates with pairwise bouts, often resulting in maiming, dismemberment, and death. These data suggest an important role for peripheral olfactory sensitivity and odor coding in the allocation of tasks in eusocial ants and support a model in which minors are multipotential olfactory generalists and majors are a hyper-aggressive soldier caste characterized by both morphological adaptions as well as a highly-specialized olfactory system focused on the detection of CHCs and other chemical cues associated with enemy non-nestmates.

## Results

### Task Allocation in Majors and Minors

To track the behavioral repertoire of adult *C. floridanus* major and minor workers within the colony, cohorts of newly eclosed, callow workers were marked with a unique paint color code on the head, thorax, and gaster. By convention, ants having eclosed within the previous 24 h were marked on day 0. The behavior of painted workers was subsequently recorded as they were observed engaging in distinctive social tasks, such as nursing (carrying eggs or larvae), cleaning (carrying trash or dead nestmates), trophallaxis, and foraging (eating crickets or Bhatkar diet or drinking sugar water). As in similar observations across many eusocial Hymenopteran, including previous studies with *C. floridanus* (Wilson 1976; Tripet and Nonacs 2004; Seeley 1982; Kolmes and Sommeijer 1984; Jeanne, Williams, and Yandell 1992), minor workers exhibited an age-associated transition from nursing (mean ± SEM = 20.83 ± 2.47 days old (d), N = 53) to foraging (51.70 ± 3.44d, N = 66) (Figure 1A). Minors performing trophallaxis with nestmates were of intermediate age (31.09 ± 4.77d, N = 22), whereas comparatively few cleaners (33.14 ± 7.53d, N = 7) were observed (Dataset S1). It should be noted that this deficit may well be an artifact of our ant husbandry protocol, which regularly incorporates manually cleaning of each colony, involving removal of trash and other debris several times per week. The earliest time point at which a minor worker was observed to forage was day 6, and nursing primarily occurred prior to day 51. In between these two time points, however, minor workers could be observed nursing or foraging. Therefore, at any one time within a colony, there may be a subset of minor worker nurses that are older than a subset of foragers, and vice versa, suggesting behavioral plasticity and task switching (Gordon 2015).

**Figure 1.**
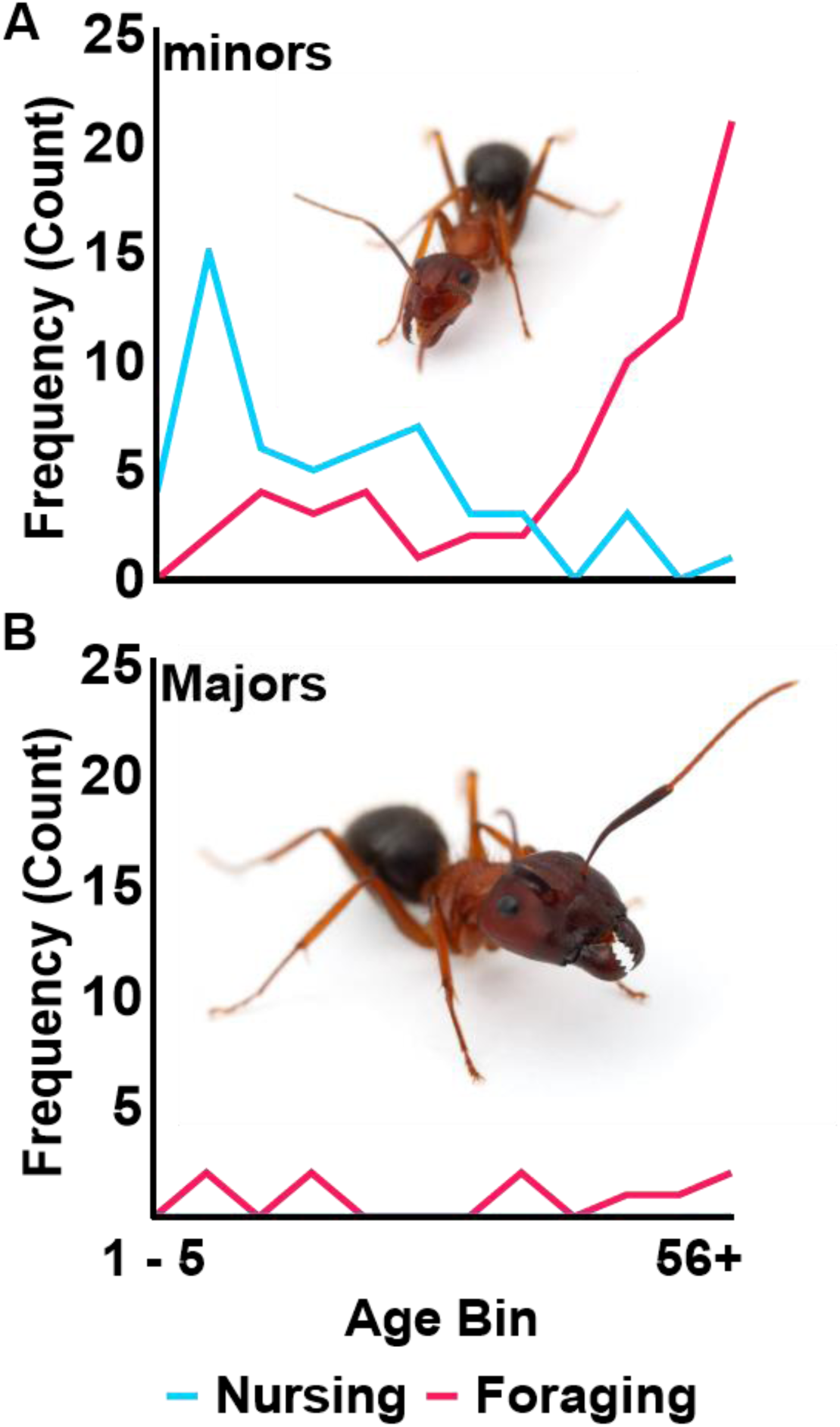
Engagement in social behavior varies between adult *C. floridanus* minor and major workers. (**A**) Minors performed the majority of tasks within the colony with an age-associated transition from nursing (N=53) to foraging (N=66). (**B**) In contrast, majors never engaged in nursing and only occasionally foraged (N=10).

In contrast to the relatively active minors, their morphologically larger major sisters only occasionally engaged in foraging (48.30 ± 14.25d, N = 10) and were never observed nursing (Figure 1B). Similarly, only a single major was ever observed cleaning (11.00, N = 1) (Dataset S1). Majors that engaged in trophallaxis (25.80 ± 6.52d, N = 5) and foraging were of a similar age to their minor counterparts (Dataset S1). This is consistent with previous studies suggesting that, within a *C. floridanus* colony, minor workers perform nearly all foraging, and that this caste-associated distinction in foraging may be regulated, at least in part, by epigenetic modifications such as histone acetylation (Simola et al. 2016).

### Peripheral Electrophysiology via the Electroantennogram

#### Positive Control

Despite the relatively impressive size of their antennae and the density of sensilla and sensory neurons (Nakanishi A 2009; Takeichi et al. 2018), electroantennogram (EAG) recordings of *C. floridanus* and other ant species routinely yield relatively low-amplitude signals compared with other insects such as *Drosophila* which have comparatively smaller antennae (Alcorta 1991; Haak 1996; Guan et al. 2014; Ferguson et al. 2020). To increase the signal-to-noise of our *C. floridanus* EAG recordings, our setup significantly restricted worker movement. Furthermore, we elected to use the heterocyclic compound 5,6,7,8-tetrahydroquinoline (TETQ), which elicits strong responses in both minors and majors, as a positive control prior to all EAG recordings in which a response threshold of at least 1.5x solvent (diethyl ether) was required to ensure suitable and consistent electrophysiological preparations.

#### General Odorants

The responses of major and minor workers were examined after stimulation with a large panel of 406 general odorants across 10 different chemical classes that were presented as 36 distinct stimulus blends (Dataset S2). Normalizing these EAG responses against solvent alone not only uncovered significant differences among odor blends between minors and majors but also revealed profound odor coding distinctions (Figure 2; Figure S1). To begin with, the olfactory responses of minors were primarily excitatory while majors displayed significantly more sub-solvent responses to odor blends than minors (Fisher’s Exact Test, P < 0.0001) (Figure 2A-B; Figure S1). These data are consistent with previous RNA sequencing studies, indicating that most olfactory gene transcripts are enriched in mature *C. floridanus* minors compared to majors (Figure S2) (Zhou et al. 2012). The diminished olfactory responses displayed by majors should not necessarily be considered anosmic, i.e., null (= 0) responses that would be comparable to the solvent-alone background controls. Rather, we interpret these large, sub-solvent responses as inhibitory and, in that context, biologically salient signals in their own right (Cao et al. 2017).

**Figure 2.**
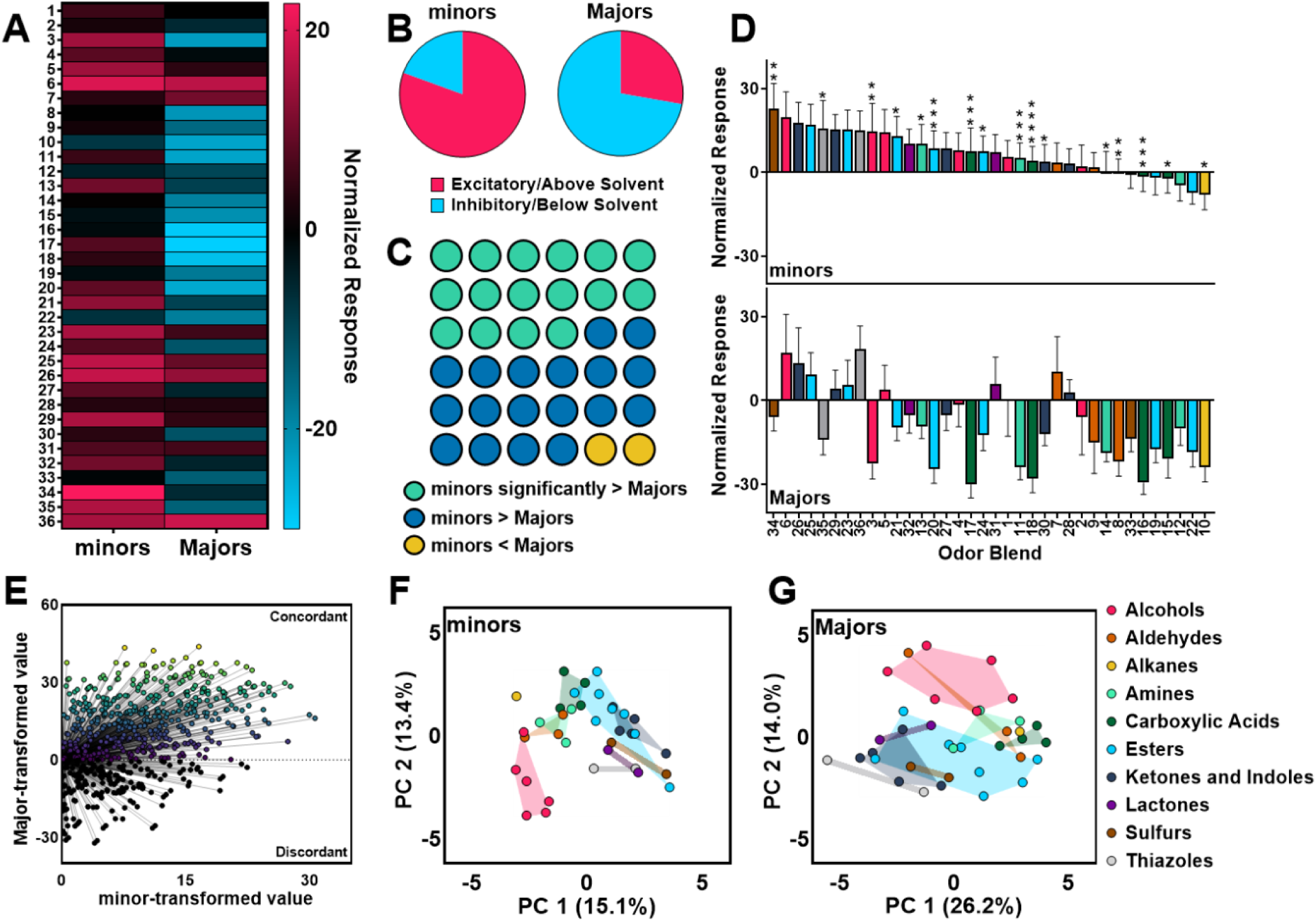
Profound differences in the olfactory response profiles of minor and major workers with respect to general odorant blends. (**A**) Heat map displaying the solvent-normalized EAG responses to the 36 odor blends in minors (N=25) and majors (N = 30). (**B**) Pie charts showing the proportion of odor blends that elicited excitatory (above solvent) responses in red and inhibitory (below solvent) responses in blue. (**C**) A dot plot indicating the odor blends that elicited a significantly higher response in minors than majors (green) as well as non-significant differences that were nevertheless higher in minors (blue) or higher in majors (yellow). No odor blends tested were significantly higher in majors than in minors. (**D**) A bar graph organized by the magnitude of responses to each odor blend from high to low in minors and the corresponding responses in majors displayed below. Colors represent chemical class as depicted in 2G. Asterisks indicate responses that were significantly different between minors and majors (Welch’s t-test, P-value: * < 0.05, ** < 0.01, *** < 0.001,. **** < 0.0001). Error bars represent S.E.M. (**E**) A visualization of Kendall’s tau whereby all 630 possible pairwise comparisons are represented with concordant pairs possessing a positive slope and discordant pairs possessing a negative slope. (**F-G**) PCA of the 10 chemical class groupings (organized by color) with convex hulls shown by the shaded regions.

A total of 16 of the 36 general odorant blends tested elicited a significantly higher response in *C. floridanus* minors than in majors (Figure 2C-D). Notably, these differences were consistent across different chemical classes, including alcohols (Blend 3), aldehydes (8), alkanes (10), amines (11, 13, 14), carboxylic acids (15-18), esters (20, 21, 24), ketones and indoles (30), sulfides (34), and thiazoles (35). The significantly higher responses to a broad range of chemical classes suggest these trends do not reflect a bias toward a particular set of odors but instead reveal a generalized increase in chemosensory sensitivity. The only exception were lactone-based odorants, which did not elicit significantly different responses between minors and majors. Of the remaining 20 odor blends, 18 elicited modestly higher responses in minors while only 2 provoked higher EAG responses in majors (Figure 2C).

Beyond uncovering these broadly attenuated responses in major workers, we also examined the extent to which minors and majors align with respect to the rank order of odors based on the response magnitude of each blend. Such information reflects the structured relationship of one odor blend to another and may align with behavioral valence. For example, the response to Blend 6 was higher than that to Blend 26 which, in turn, was higher than to Blend 25 in both minors and majors (i.e., concordance; Figure 2D). In contrast, the response to Blend 34 was higher than that to Blend 6 in minors, but the opposite was true for majors (i.e., discordance; Figure 2D). This analysis revealed that, despite profound differences in olfactory sensitivity, for any given pair of odor blends minors and majors tend to assign the same rank order (τ = 0.427, P < 0,001; Figure 2E). Among the 630 possible pairwise comparisons, 449 were concordant and 180 were discordant with one tie value (Figure 2E). These data suggest that, while response magnitude may vary significantly, the rank relationships between pairs of odor blends are remarkably consistent between worker castes (Figure 2D-E). At the same time, this analysis also highlights a large number of discordant rankings that serves to further distinguish the physiological response profiles of these morphological and behaviorally distinct castes (Figure 2E).

Principle component analysis (PCA) illustrates the similarities and differences of minor and major olfactory responses with respect to stimuli chemical class (Figure 2F-G). Here, alcohols formed a cluster apart from most other chemical groups in both minors and majors, which suggests these chemicals are relatively distinct insofar as their ability to be discriminated apart from other stimuli. Esters, on the other hand, partially overlapped with ketones and indoles and were generally more dispersed, likely owing to the range of both excitatory and inhibitory responses within this class. Across all chemical classes, however, tighter clusters were formed in minors (Figure 2F), reflecting their broad and consistent excitatory responses to most odor blends (Figure 2A-D). In contrast, a greater degree of overlap among chemical classes was observed in majors (Figure 2G), where odor blends elicited a greater range of primarily inhibitory responses (Figure 2A-D). Taken together, these data further support the hypothesis that olfactory sensitivity is linked to caste identity and task allocation between minor and major castes in *C. floridanus*.

#### Trail Pheromone

Extending these studies beyond general odorants to pheromones and other chemical blends that are likely to be important sources of social information, we next focused on the trail pheromone nerolic acid. Previous studies have demonstrated this to be produced in the rectal bladder of *C. floridanus* workers and elicit a strong trail-following phenotype that is not evoked by its double-bond structural isomer geranic acid (Haak 1996). Because purified nerolic acid is difficult and expensive to obtain, the lack of geranic acid responses facilitated employing a commercially available isomeric mixture (Sigma-Aldrich, CAS 459-80-3) that accordingly is hereafter denoted as nerolic acid. In EAGs, minors displayed significantly higher and dose-dependent responses at 10^−5^ M (Welch’s t-test, P < 0.01) and 10^−3^ M (P < 0.05) nerolic acid than majors whose responses were slightly below solvent-alone levels at all concentrations (Figure 3A). These peripheral responses are consistent with the reduced level of foraging activity observed in majors (Figure 1B). The behavioral responses of minor and major workers to nerolic acid were also investigated using a trail-following bioassay (Figure 3B). In solvent-alone controls, both minors and majors preferred to aggregate near the corners and edges of the foraging arena (Figure 3C-D). Importantly, only minors followed the nerolic acid trail pheromone cue (Figure 3E) to which *C. floridanus* majors seem indifferent (Figure 3F). Indeed, the majority of minors aligned themselves along the nerolic acid trail and followed it throughout its length, whereas majors did not follow the trail and instead aggregated closer to the arena edges. Taken together, these observations suggest that the allocation of foraging to the colony’s minor workers may be regulated, at least in part, by differential peripheral olfactory sensitivity to a discrete subset of salient cues that includes nerolic acid.

**Figure 3.**
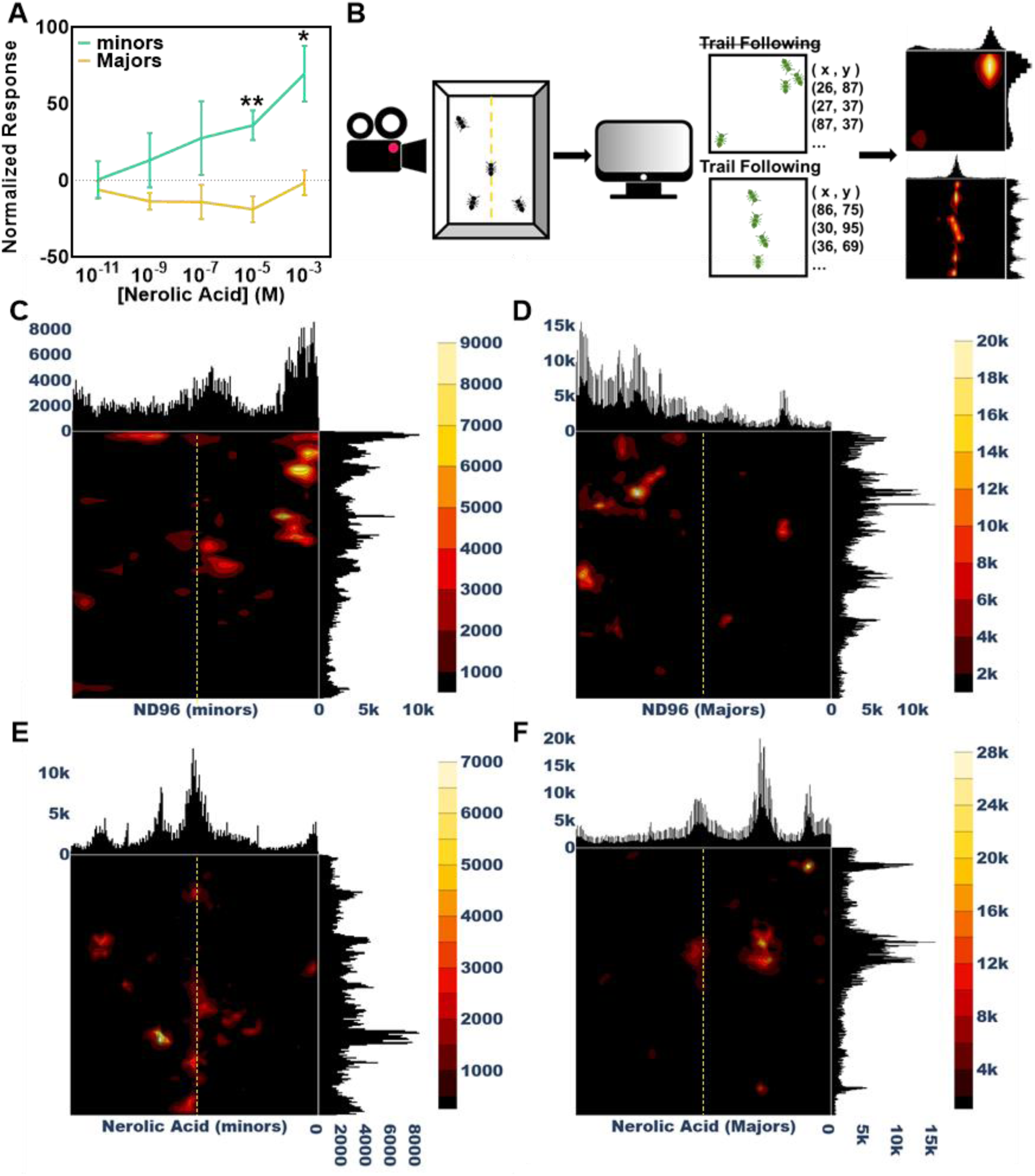
High olfactory sensitivity and trail following in response to nerolic acid in minors but not majors. (**A**) Minors, but not majors, exhibited robust, dose-dependent excitatory responses to nerolic acid and displayed significantly higher responses at 10-5 M (N = 5) (Welch’s t-test, P-value: ** = 0.002859) and 10-3 M (P-value: * = 0.013602). Error bars represent S.E.M. (**B**) Schematic of the foraging bioassay. Histograms correspond to the x,y coordinates of ant bodies detected throughout the videos as a percentage of the width and height of the foraging arena. The heat maps are density contour plots with darker regions indicating areas of low density where ants were less likely to be found and brighter regions indicating areas of high density where ants were more likely to be found. (**C-D**) Neither minors nor majors followed the ND96 solvent alone. (**E-F**) Minors followed artificial nerolic acid trails whereas majors did not. Chemical trails are denoted as a yellow dashed line.

#### Cuticular Extracts

We next compared major and minor worker responses to CHCs and other hydrophobic cuticle components by focusing on non-nestmate cuticle extracts, which are acutely evocative stimuli in the context of aggression (Ferguson et al. 2020). Here, whole-body cuticular extracts of adult minor workers were obtained using a hexane soak at dilutions equivalent to the surface content of 0.001, 0.05, 0.25, 0.50, 0.75, or 1 individual ant. These cuticular extracts uniformly elicited robust and generally dose-dependent excitatory EAG responses from both minors and majors (Figure 4A). While the magnitude of responses was similar in both minors and majors across the highest extract concentrations, majors displayed significantly higher excitatory responses than minors at the lowest concentration tested (equivalent to 0.1% of an ant; Welch’s t-test, P < 0.05). Importantly, the robust excitatory responses in majors elicited by CHC extracts contrast with the observed broad inhibitory responses to general odorant blends and trail pheromone (Figures 2A-D, 3A). These provocative electrophysiological characteristics add to the distinction between *C. floridanus* minors and major worker castes.

**Figure 4.**
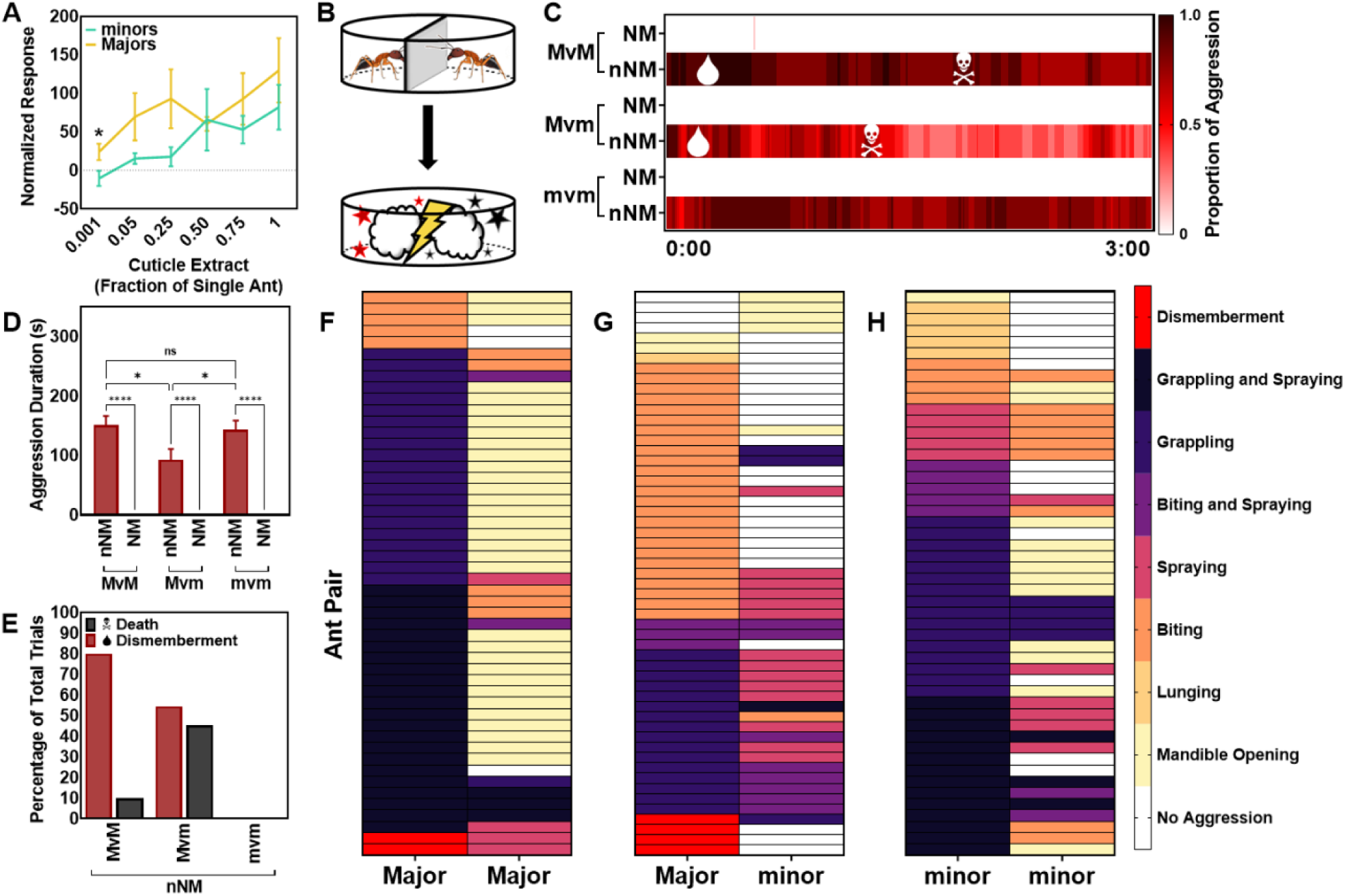
High olfactory sensitivity to cuticle extract in minors and majors. (**A**) Both minors and majors exhibited robust, dose-dependent excitatory responses to cuticle extract from non-nestmate minor workers. Majors were significantly more responsive to extract at the lowest concentration tested compared with minors (N = 5) (Welch’s t-test, P-value: * = 0.046383). Error bars represent S.E.M. Minors and majors both accepted nestmates and aggressed non-nestmates, but only majors dismembered and killed their opponents. (**B**) Schematic of the bioassay depicting the acclimation period (top) and aggression bioassay (bottom). (**C**) A heat map showing the elapsed time of the 3-min aggression bioassay along the X-axis. The heat map represents the proportion of trials where aggression was observed at each timepoint (N=10-11). The mean time at which dismemberment (droplet) or death (skull and crossbones) occurred is indicated along the heat map. Notably, minor vs. minor bouts never results in dismemberment or death. (**D**) The mean aggression duration (in seconds) between nestmates and non-nestmates (N = 10-11) (Two-Way ANOVA with Tukey’s correction, P-value: ns = not significant, * < 0.05, **** < 0.0001). (**E**) The percentage of trials that results in at least one instance of dismemberment or death. (**F-H**) A heat map with pairs of combative non-nestmate ants, with each individual ant represented as a cell within a row. The color of the cell indicates their aggressive behavior (mandible opening, lunging, biting, spraying, biting and spraying, grappling, grappling and spraying, and dismemberment) at a randomly sampled time point throughout the video bioassay trial (N = 50 – 55).

To further examine whether morphologically distinct major workers do indeed represent a dedicated and highly specialized soldier caste within the colony, we next quantified several elements of the aggressive behaviors characteristically seen in interactions between pairs of minors and majors using a non-nestmate recognition aggression bioassay (Figure 4B). While both minors and majors accurately accepted nestmates and, in contrast, dramatically aggressed non-nestmates (Figure 4C-D), particularly aggressive interactions that rapidly resulted in incapacitation or death through dismemberment of appendages were only observed in bouts involving majors (Figure 4C, E-H).

Indeed, majors were responsible for all acts of dismemberment and death and were profoundly adept at rapidly killing smaller non-nestmate minor worker opponents, which resulted in significantly shorter fights than in major vs. major and minor vs. minor bouts (Tukey’s comparison, P < 0.0001; Figure 4D). There were also notable differences in the fighting strategy of majors and minors, which varied depending on which caste they were fighting (Figure 4F-H). In pairwise bouts between two majors, there was often a tactically superior fighter that would dominate their opponent by grasping them with their mandibles in a prolonged grapple that was occasionally accompanied by sprays of formic acid and dismemberment (Figure 4F). This grapple restrained and inhibited offensive attacks from opponents. Majors restrained in a grapple responded with mandible opening but nevertheless only rarely launched a successful counterattack of biting, spraying, or grappling. When facing smaller minor opponents, majors engaged in more biting and less grappling (Figure 4G). This is likely due to the efficiency of their attacks, which often led to dismemberment and death. Minors were less likely to attack majors and instead would rely on spraying formic acid to potentially disorient them and thereby escape devastating lethal attacks from majors. Fights between minors were more dynamic but less lethal; these attacks never resulted in dismemberment (Figure 4H). While some individuals occasionally secured a dominant grappling position, minors were better able to respond with biting, spraying formic acid, and grappling (Figure 4H) than the victims in major vs. major fights (Figure 4F). Nevertheless, even when majors were dominated by their opponent’s grapple, they continued to display aggression through mandible opening (Figure 4F). In contrast, when many minor workers were victims of aggression they did not attempt to fight back (Figure 4G-H). Taken together with the electrophysiology, these behavioral distinctions suggest that non-nestmate recognition signals present on the cuticle are detected, perceived, and acted on by both majors and minors. However, majors are more aggressive and therefore vastly superior fighters that are capable of rapidly killing their opponents.

## Discussion

In eusocial ant colonies, behavioral patterns such as nursing, foraging, and colony defense emerge from the collective behavior of workers. These emergent social behaviors occur through a decentralized distribution of work across members of the colony known as task allocation (Gordon 1996), which is the outcome of intrinsic and extrinsic factors (Gordon 1996; Crall et al. 2018; Gordon, Dektar, and Pinter-Wollman 2013; Cole et al. 2010). While behavioral caste is often associated with age (Seeley 1982; Kolmes and Sommeijer 1984; Wilson 1976), ant workers also switch tasks in response to a wide range of chemical and tactile signals that convey situational changes in colony requirements (Crall et al. 2018; Gordon 1989). These peripheral stimuli are integrated and decoded in various parts of the ant brain, which has a remarkable capacity to discriminate subtle features of information (Najar-Rodriguez et al. 2010; Silbering et al. 2008). Here, we have extended these studies through a large electrophysiological screen of general odorants, pheromones, and cuticular odor blends complemented by a series of behavior bioassays to test the hypothesis that variation in peripheral olfactory sensitivity of *C. floridanus* major and minor workers correlates with caste-specific differences in their behavioral repertoire.

Consistent with previous studies in *C. floridanus* (Simola et al. 2016) and other ant species (Wilson 1984), we found that minor workers performed the majority of tasks within the colony, such as tending to the brood and foraging for food (Figure 1). These responses aligned with the ability of minors to robustly detect and follow nerolic acid as a trail pheromone (Figure 3). Major workers were, by comparison, less active and displayed a diminished behavioral repertoire insofar as they were never involved in brood care, displayed only low levels of foraging and food exchange through trophallaxis, and did not follow trail pheromone (Figure 1, Figure 3, Dataset S1). This apparent ‘laziness’ explains why the precise role of *C. floridanus* majors has remained somewhat speculative.

When examining these behavioral characteristics through the prism of peripheral olfactory physiology, profound differences in sensitivity and odor coding were identified between *C. floridanus* minors and majors. To begin with minor workers were significantly more sensitive to general odorants (Figure 2C) and trail pheromone (Figure 3A). Furthermore, general odorant stimuli elicited primarily excitatory responses from minors whereas majors displayed primarily sub-solvent, inhibitory responses (Figure 2). That the expanded repertoire and higher engagement in social behavior by minor workers is associated with broad excitation and detection of general odorants and pheromones and contrasts with low activity and inhibitory responsiveness in majors provides an intriguing link between the regulation of peripheral olfactory physiology and task allocation. While we are as yet unable to fully appreciate the odor coding implications of excitatory versus inhibitory EAG responses, these data align with models of task allocation that propose behavioral performance depends on internal differences between individuals arising from “hard-wired” stimulus thresholds that determine the probability of engaging in a task (Beshers, Robinson, and Mittenthal 1999).

Interestingly, in contrast to general odorants and trail pheromone, both minors and majors displayed robust, dose-dependent excitatory responses to non-nestmate cuticular extracts (Figure 4). Importantly, the olfactory responses of majors to these stimuli were slightly higher than those of minors at most concentrations tested and significantly higher at the lowest concentration tested (the equivalent of 0.1% of an ant, Figure 4). If task performance depends on physiological stimulus thresholds, then it stands to reason that *C. floridanus* majors are likely to be involved in robust aggressive behaviors towards non-nestmates to the exclusion of other tasks. Consistent with this hypothesis, we found that while both minors and majors could accurately discriminate between nestmates and non-nestmates, major workers were significantly more effective at rapidly dismembering and killing non-nestmates during aggressive interactions. Moreover, they were relentlessly aggressive even when facing a tactically superior opponent (Figure 4). This distinctive electrophysiology and aggressive behavior suggest that while non-nestmate recognition signals are detected, perceived, and acted on by both castes, majors are categorically and quantitatively more effective fighters, owing not only to their superior size and mandibular characteristics but also and importantly due to their enhanced and focused chemosensory responses to discrete non-nestmate stimuli. Indeed, the olfactory system of majors seems to be selectively specialized for the detection of CHCs and other salient compounds linked to a narrow range of social behaviors associated with the recognition and aggressive rejection of non-nestmates as potential threats to colony integrity. We posit that these profound differences in olfactory sensitivity are evolutionary adaptations that reflects the unique physiology and behavioral specialization of each morphological caste.

Our data are consistent with the hypothesis that minors are multipotential jacks-of-all trades, engaged in a diverse set of social behaviors in the colony with broad, excitatory olfactory sensitivity to general odorants and pheromones. Majors, however, are a highly specialized soldier caste dedicated to colony defense, the olfactory system of which is fine tuned to detect non-nestmate chemical signatures. Taken together, these results suggest that directed shifts in olfactory sensitivity play important roles in establishing and maintaining caste identity as well as the allocation of social behaviors within a eusocial collective.

## Materials and Methods

### Animal husbandry

Six laboratory-reared colonies of *Camponotus floridanus* (Buckley 1866) originating from field collections from the Sugarloaf Key (D601) and the Fiesta Key (C6, K17, K19, K34 and K39) in South Florida, USA, were generously gifted by the laboratories of Dr. J. Liebig (Arizona State University) and Dr. S. Berger (University of Pennsylvania), respectively. The colonies were separately maintained at 25°C with an ambient humidity of approximately 70%. Colonies were stored in an incubation chamber with a 12 h light:12 h dark photoperiod. Each colony was provided with Bhatkar diet, crickets, 10% sucrose solution, and distilled water three times per week. Adult minor and major workers were used for all experiments.

### Paint marking, behavioral monitoring, and collections

Callow workers were identified based on their soft, light-colored cuticle, low mobility, and proximity to the brood pile. Ants with these characteristics were likely to be less than 24 h post-eclosion at the time of collection. These callow workers were briefly anesthetized with CO2, and Sharpie oil-based paint pens were used to mark the head, thorax, and gaster with a unique color code. Prior to making behavioral observations, colonies were removed from the incubation chamber and allowed to acclimate after handling for at least 5 minutes. Observations were made between ZT0–ZT12 and lasted approximately 15 minutes. If the colony was disturbed, for example when removing trash or replacing food, no observations were made that day. The behavior of age-known ants engaged in pre-specified behaviors (carrying eggs, carrying larvae, performing trophallaxis, eating crickets, eating Bhatkar, and drinking sugar water) that could be identified based on their paint code was then recorded as observed. To collect ants for downstream electroantennography and behavioral bioassay analyses, individual ants were randomly sampled from among the different colonies tested.

### Electroantennography

Electroantennograms (EAGs) were conducted using an IDAC-232 controller (Ockenfels Syntech GmbH, Buchenbach, Germany) and data were initially collected and stored on EAG2000 software (Ockenfels Syntech GmbH). Odorants were delivered using a Syntech CS-05 stimulus flow controller (flow rate of 1.5 cm^3^ s^−1^; Ockenfels Syntech GmbH). Minors were placed in a 20-μl disposable pipette tip that was modified such that the tip opening was sufficiently wide to allow the unimpeded exposure of the head and antennae. Majors were placed in modified 200-μl pipette tips to accommodate their wider head. To prevent unwanted movement from the ant that might otherwise interfere with the quality of the recording, the head and mandibles of the ant were restricted with wax in addition to the right antennae. Borosilicate glass capillaries (FIL, o.d. 1.0 mm, World Precision Instruments, Inc.) were prepared using a P-2000 laser micro-pipette puller (Sutter Instruments). Both the reference electrode and the recording electrode were backfilled with 10^−1^ mol l^−1^ KCl and 0.05% PVP buffer and placed over tungsten electrodes. Due to the armor-like exoskeleton of the ant, a 30-gauge needle was required to puncture the right eye prior to inserting the reference electrode. The recording electrode was placed over the distal tip of the left antenna. The resulting signals were amplified 10× and imported into a Windows PC via an intelligent data acquisition controller (IDAC, Syntech, Hilversum, The Netherlands) interface box. These recordings were analyzed using EAG software (EAG Version 2.7, Syntech, Hilversum, The Netherlands) such that stimulus-evoked response amplitudes were normalized using linear interpolation and then subtracting the response amplitude of control (solvent alone) responses.

The TETQ positive control was diluted in diethyl ether at a concentration of 10^−1^ M. Odor cartridges were prepared with 10 μl of solution. Prior to all electrophysiological recordings, the ant was first exposed to diethyl ether, then TETQ, and then another exposure to diethyl ether. Normalization of the TETQ response was then accomplished through linear interpolation vis-à-vis EAG2000. If the response was greater than 1.5x solvent, then the experiment would commence. Otherwise, the experimental recording was not conducted. Odor blends were diluted in ND96 buffer. Each odorant was diluted to a concentration of 10^−3^ M. Odor cartridges were again filled with 10 μl of solution. Recordings were conducted in the following order: ND96, odor blends 1-18, ND96, odor blends 19-36, ND96. In this way, responses could be normalized to solvent responses recorded across the duration of the trial to account for antennae degradation over time throughout the assay.

Cuticular extracts were obtained by soaking 40 minor nestmates in 8 ml hexane for 30 min. This cuticle soak was then decanted, and the hexane was evaporated using compressed nitrogen. The remaining contents of the extraction were then resuspended in hexane so that odor cartridges were filled with 20 μl hexane or hexane plus extract at the appropriate concentration (equivalent to cuticle soak obtained from 0.001, 0.05, 0.25, 0.50, 0.75, or 1 whole ant). When testing odor responses, a handheld butane torch (BernzOmatic, Worthington Industries, Columbus, OH, USA) was used to volatilize the cuticular compounds by heating the odor cartridge for 1.5 s. Odors were introduced in the following order: hexane, cuticle extract, hexane. Given that nerolic acid is detected and elicits robust trail following in *C. floridanus* workers but its isomer geranic acid is neither detected nor followed (Haak 1996), nerolic acid was obtained as a mixture of isomers (Sigma-Aldrich, CAS 459-80-3) comprised of roughly 70% trans-geranic acid and about 20-25% nerolic acid (personal communication with manufacturer). The nerolic acid was then serially diluted to concentrations of approximately 10^−11^, 10^−9^, 10^−7^, 10^−5^, and 10^−3^ M. Odors were then introduced in the following order: ND96, nerolic acid, ND96 so that, as before, responses could be normalized to solvent responses recorded across the duration of the trial to account for antennae degradation over time throughout the assay.

### Kendall’s Tau Visualization

To visualize the results from Kendall’s rank correlation coefficient (Tau-b, τ), we used a novel transformation of our solvent-normalized EAG response data. We began by plotting minors’ responses to each general odorant blend along the x-axis and majors’ responses along the y-axis. We then examined every possible pairwise comparison between blends (630 in total) moving from left to right along the Cartesian plane. The coordinate on the left was then set to (0,0) and its corresponding pair transformed accordingly. For example, a pair of coordinates (10,15) and (20,40) would become (0,0) and (10,25). In this manner, all 630 comparisons were normalized to the x,y intersect at (0,0) with concordant pairs presented as lines with a positive slope and discordant pairs presented as lines with a negative slope.

### Principal Component Analysis (PCA)

PCA were performed using ClustVis (Metsalu and Vilo 2015), and the data was exported to Prism (Graphpad Software ©) to create the PCA plots. Convex hulls were added manually.

### Behavioral Bioassays

#### Trail-Following Bioassay

Foraging arenas were constructed using a sheet of Whatman with 70 μlof either solvent (ND96) or nerolic acid (as a mixture of isomers) distributed along the length of the center of the paper using a pipette. For each replicate, ten adult minor or major workers were removed from a randomly selected colony and placed in a Petri dish to acclimate for at least 5 min. After allowing the solvent to evaporate for 5 min, the ants were introduced into the arena and their activity was subsequently digitally recorded for 15 min. To identify individual ants in the digital video recordings of each foraging bioassay, videos were first cropped using Adobe Premiere Pro to include only the foraging arena of the bioassay, and the brightness and contrast were adjusted so that the background appeared white and the ants appeared black. This video was then exported as a series of TIFF files at a frame rate of 1 frame per second. The OpenCV and Numpy libraries in Python were then used to convert the TIFF files to grayscale and identify the x,y coordinates of all pixels with a grayscale value less than 100 (i.e., gray to black) as a percentage of the total foraging arena which would correspond to the location of the ants. Plotly, together with Pandas, was then used to generate 2D histogram contour plots.

#### Aggressive Recognition of non-Nestmates

Aggression bioassays were modelled after a previously published method (see (Ferguson et al. 2020)). Briefly, nestmate and non-nestmate workers were randomly assigned to the following treatment groups: minor vs. minor, minor vs. major, and major vs. major. Two workers were placed in each half of an individual well in a six-well culture plate with a plastic divider separating the ants in the middle. The ants were allowed to acclimate for 13 min before removing the divider and recording their interactions for 3 min using a digital, high-definition camera (Panasonic® HC-V750). Videos were then scored by a blinded observer. Each frame of the video received a binary score of either 0 (no aggression) or 1 (aggression). Aggression behavior included mandible opening, lunging, biting, spraying formic acid, grappling, and dismemberment. To determine the combat style of majors and minors, we randomly sampled five different time points in which aggression was observed between non-nestmates from each video. This resulted in 50 observations for majors vs. majors and for minors vs. minors, and 55 observations for majors vs. minors. The behavior of each ant was then recorded.

In certain circumstances, only one ant was aggressive. This was due to factors such as avoidance behaviors such as running away and incapacitation. In these instances, the ant received a score of “No Aggression” because they were not actively engaged in an aggressive act.

#### Statistical Analyses

Prism (Graphpad Software ©) was used to create all graphics figures and perform all statistical testing. The exception is the 2D histogram contour plot for the trail following bioassay, which relied on the Python library Plotly. A summary of the raw data for the figures and statistical analyses presented in this manuscript can be found in Datasets S1-13.

## Acknowledgments

We thank the laboratories of our colleagues Dr. J. Liebig (Arizona State University) and Dr. SL Berger (University of Pennsylvania) for ant collections. We also thank Drs. D.P Abbot, MJ Greene and HW Honegger for their thoughtful insights on the manuscript. Lastly, we thank members of the Zwiebel lab for their suggestions throughout the course of this work, Drs. S. Ochieng for ant rearing and technical help, and AM McAinsh for editorial assistance.

## Author contributions

Conceived experiments: STF and LJZ; Performed research: STF, IB, and NDE; Analyzed data: STF, IB, and NDE; Wrote the paper: STF, IB, NDE, and LJZ. Approved the final manuscript: STF, IB, NDE, and LJZ.

## Figures and Figure Legends

**Figure S1.**
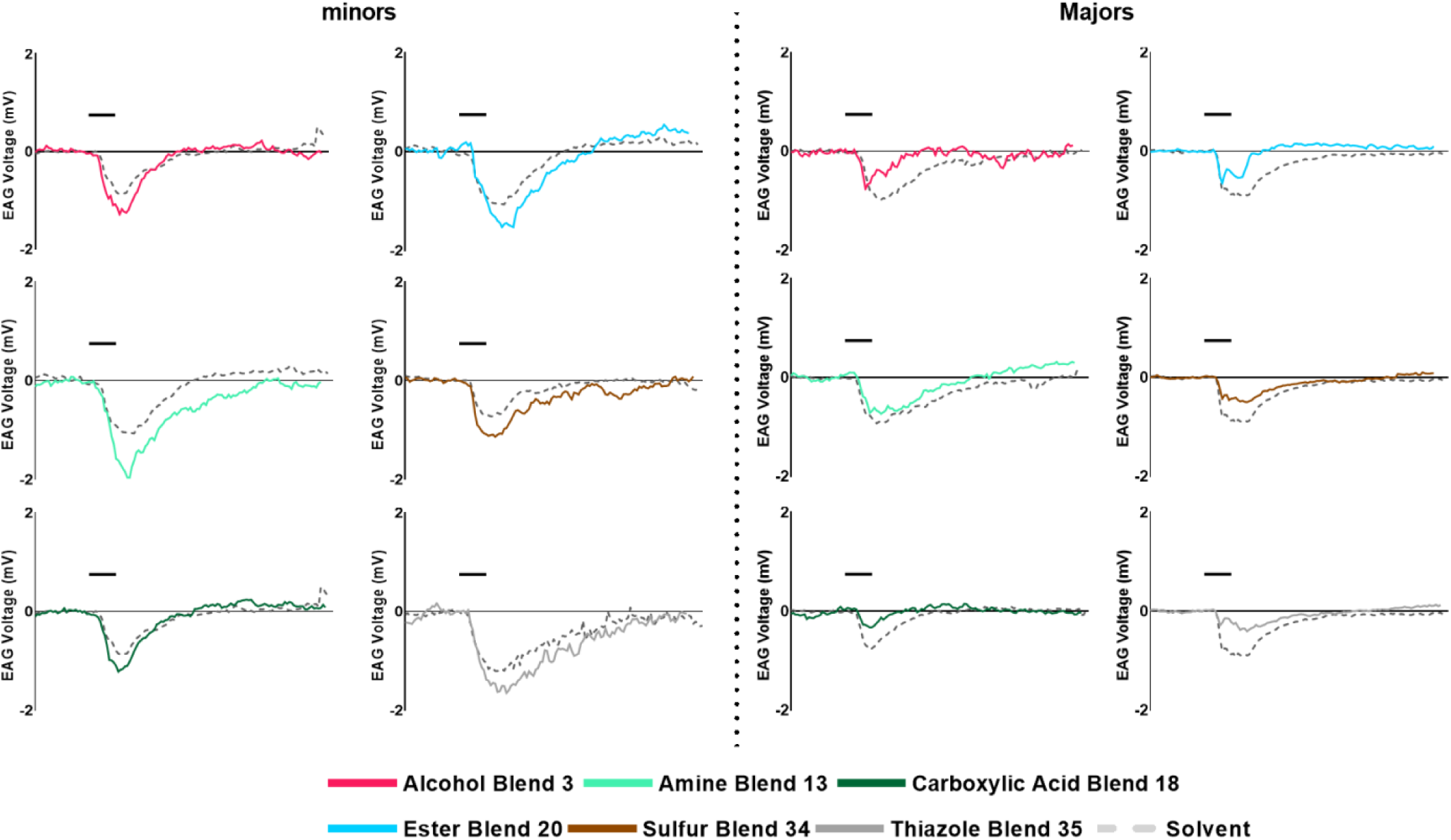
Representative EAG traces display excitatory responses in minors and inhibitory responses in majors. Representative EAG traces from six different odor blends: Alcohol Blend 3 (red), Amine Blend 13 (light green), Carboxylic Acid Blend 18 (dark green), Ester Blend 20 (blue), Sulfur Blend 34 (brown), and Thiazole Blend 35 (gray) so minors (left) and majors (right) with the average ND96 solvent response across the recording displayed by the dashed gray line.

**Figure S2.**
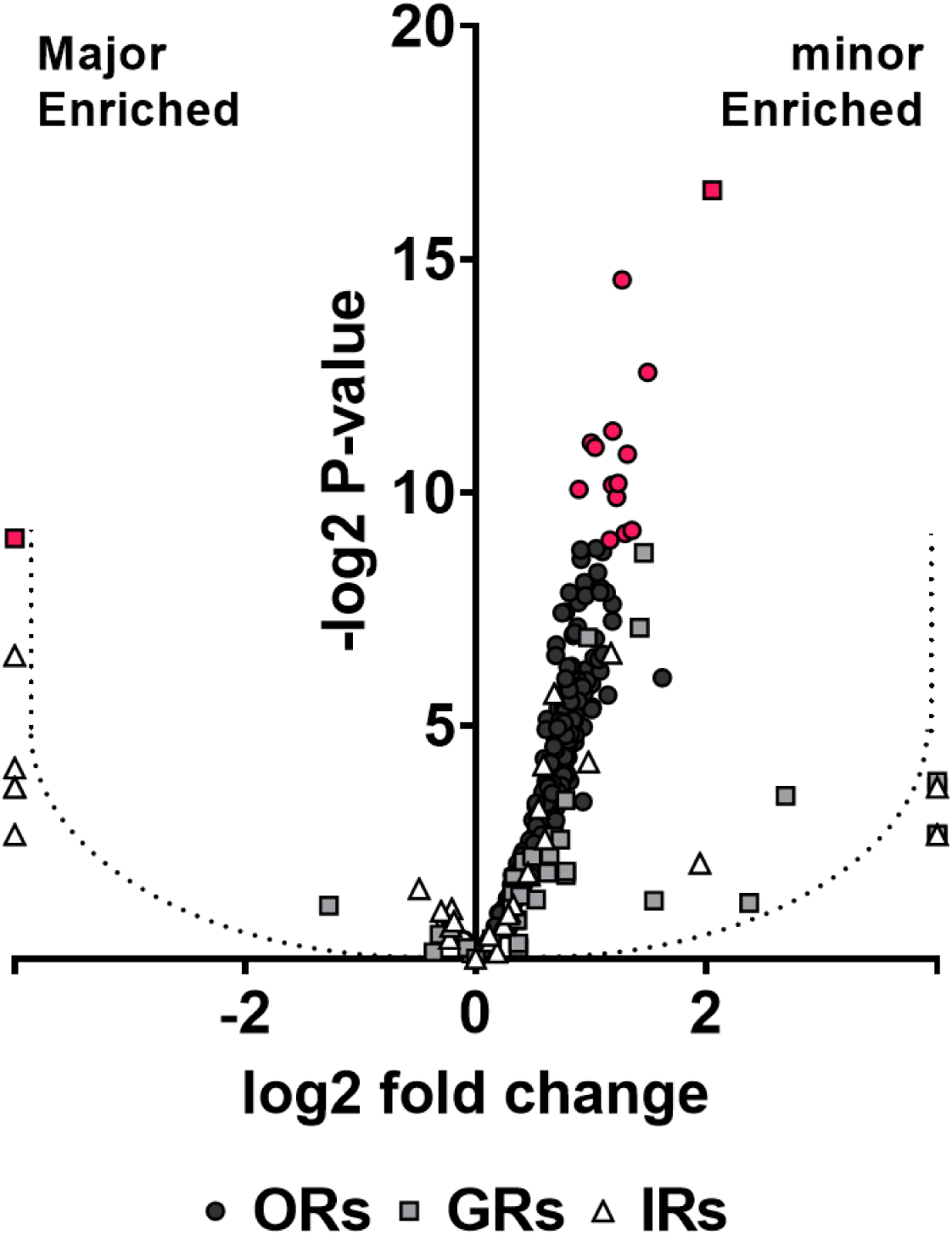
Chemoreceptor transcripts are enriched in minor workers compared with majors. The antennae of minor *C. floridanus* workers are enriched for the three primary classes of chemoreceptors: ORs, GRs, and IRs. Figure derived from data published in (Zhou et al. 2012). Red fill indicates genes that were significantly enriched in either caste. Points that fall to the left or right of the dotted lines indicate genes that were not detected in minors and majors, respectively.

## Datasets S1-13. Raw data

(Dataset S1) Behavioral tracking and observations. (Dataset S2) A list of the general odorant blends tested. (Dataset S3) EAG responses of minors and majors to the general odorant blends. (Dataset S4) PCA output. (Dataset S5) Representative EAG traces. (Dataset S6) EAG responses of minors and majors to nerolic acid. (Dataset S7) EAG responses of minors and majors to cuticular extract. (Dataset S8) Aggression bioassay scoring (0 = no aggression, 1 = aggression). (Dataset S9) Aggressive behavior between non-nestmates at randomly sampled time points. (Datasets S10-13) Output data for the trail following heatmaps for minors (ND96), majors (ND96), minors (nerolic acid), and majors (nerolic acid), respectively.

